# Tertiary Lymphoid Structures Sustain Cutaneous B cell Activity in Hidradenitis Suppurativa

**DOI:** 10.1101/2023.02.14.528504

**Authors:** Margaret M. Lowe, Jarish N. Cohen, Madison I. Moss, Sean Clancy, James Adler, Ashley Yates, Haley B. Naik, Mariela Pauli, Ian Taylor, Austin McKay, Hobart Harris, Esther Kim, Scott L. Hansen, Michael D. Rosenblum, Joshua M. Moreau

**Affiliations:** Department of Dermatology, UCSF; Bioinformatics and Genomics Master’s Program, University of Oregon; Cancer Early Detection Advanced Research Center, OHSU; TRex Bio; Department of Surgery, UCSF; Division of Oncological Sciences, OHSU; Department of Dermatology, OHSU

**Keywords:** Hidradenitis suppurativa, Acne inversa, tertiary lymphoid structures, B cells, T peripheral helper cells, spatial transcriptomics, single cell RNA sequencing

## Abstract

**Background:** Hidradenitis suppurativa (HS) skin lesions are highly inflammatory and characterized by a large immune infiltrate. While B cells and plasma cells comprise a major component of this immune milieu the biology and contribution of these cells in HS pathogenesis is unclear.

**Objective:** We aimed to investigate the dynamics and microenvironmental interactions of B cells within cutaneous HS lesions.

**Methods:** We combined histological analysis, single-cell RNA-sequencing (scRNAseq), and spatial transcriptomic profiling of HS lesions to define the tissue microenvironment relative to B cell activity within this disease.

**Results:** Our findings identify tertiary lymphoid structures (TLS) within HS lesions and describe organized interactions between T cells, B cells, antigen presenting cells and skin stroma. We find evidence that B cells within HS TLS actively undergo maturation, including participation in germinal center reactions and class switch recombination. Moreover, skin stroma and accumulating T cells are primed to support the formation of TLS and facilitate B cell recruitment during HS.

**Conclusion:** Our data definitively demonstrate the presence of TLS in lesional HS skin and point to ongoing cutaneous B cell maturation through class switch recombination and affinity maturation during disease progression in this inflamed non-lymphoid tissue.

## Introduction

Hidradenitis suppurativa (HS) is a chronic inflammatory skin condition primarily occurring in intertriginous regions of the body affecting approximately 1% of the US population^1^. HS may progress from acne-like inflammatory nodules to recurrent abscesses, formation of extensive sinus tracts, and deep-seated fibrosis and scarring^1^. Treatment strategies for HS include antimicrobial washes and antibiotics, steroids, hormonal therapies, anti-inflammatory biologics, and surgery to resect areas of advanced fibrosis. Targeted biologic treatments such as anti-TNF? agents however, do not provide clinical benefit to all patients, which may be linked to the extent of immune activation within the skin prior to treatment^1^. A more detailed understanding of the immune pathogenesis in HS is required to catalyze development of effective targeted therapies.

Immune dysregulation surrounding the hair follicle following follicular occlusion is hypothesized to initiate the aberrant inflammatory process characteristic of HS plaques; however, exact disease etiology is unclear^1–3^. For example, immune infiltrates within HS lesions include overrepresentation of cell subsets that are rarely observed in healthy skin and other inflammatory skin diseases, especially B cells and plasma cells^4–6^. In addition, antibodies recognizing a broad array of autoantigens have been identified in patient sera as well as lesional skin suggesting contribution of B lineage cells to disease pathogenesis^6–10^. Limited studies involving B cell depletion with rituximab hint that modulation of these cells may have therapeutic benefit^11,12^. Lymphoid aggregates comprising B cells and T cells have been described in HS lesions and are a recognized histological feature of this disease but it is unknown if organized tertiary lymphoid structures (TLS) capable of sustaining cutaneous B cell activity *in situ* are present^1,4,13^. To understand the dynamics and microenvironmental interactions of B cells within HS lesions, we employed histological analysis, scRNAseq, and spatial transcriptomics. Our data definitively demonstrate the presence of functional TLS in this disease and define the molecular identity and contributing stromal niche for generation of these structures in diseased HS skin.

## Results & Discussion

### Tertiary Lymphoid Structures are present in HS skin

Nine cases of HS and one case of normal skin were selected for histopathologic and immunohistochemical evaluation. HS cases were subdivided into those that showed no evidence of lymphoid aggregates (LA) or tertiary lymphoid structures (TLS) (3 cases), cases with LA but no discernable TLS (2 cases), and cases with TLS (4 cases) (Fig. 1A). On hematoxylin and eosin staining (H&E), TLS and LA were defined as small, discreet, well-circumscribed collections of mononuclear cells with and without a germinal center reaction, respectively. To provide supportive evidence of authentic TLS, tissue sections were stained for complement receptor 2, also known as CD21, to highlight follicular dendritic cells (FDCs) that are normally present in germinal centers (Fig. 1B)^14^. CD21 immunohistochemistry (IHC) demonstrated discrete collections of FDCs in HS cases with TLS and absence of positive staining in cases with only LA, those with no LA/TLS, and normal skin.

**Fig 1.**
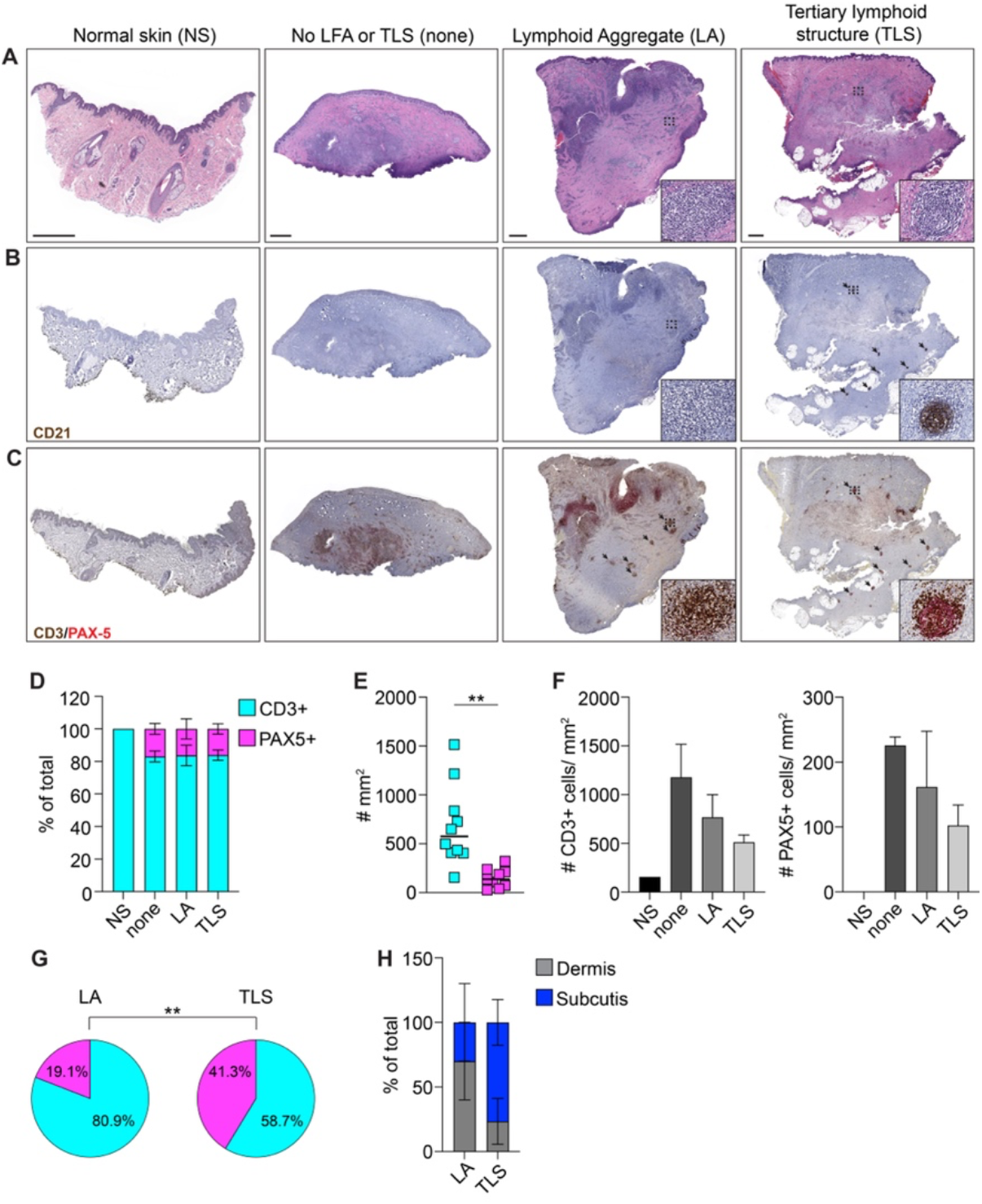
Tertiary Lymphoid Structures are present in HS skin. (A) Low power photomicrographs of hematoxylin and eosin stained sections of normal and HS skin with the indicated features. Dashed boxes indicate representative lymphoid aggregate (LA) or tertiary lymphoid structure (TLS) and insets show a higher power image within dashed boxes; bar= 2 mm. (B) Low and higher power (insets) photomicrographs of CD21 immunohistochemistry (IHC) on normal and HS skin. (C) Low and higher power (insets) photomicrographs of multiplex CD3 (brown chromogen)/ PAX5 (red chromogen) IHC on normal and HS skin. (D) Frequency of CD3+PAX5- and CD3-PAX5+ cells in normal skin and each subset of HS skin based on presence or absence of LA or TLS. (E) Number of CD3+PAX5- and CD3-PAX5+ cells in HS skin per unit area. (F) Number of CD3+PAX5- and CD3-PAX5+ cells in each subset of HS skin based on presence or absence of LA or TLS. (G) Frequency of CD3+PAX5- and CD3-PAX5+ cells in LA or TLS annotated regions. (H) Frequency of the microanatomic distribution in the dermis or subcutis of LA and TLS in HS skin. **p<0.01.

To evaluate the distribution of the lymphocytes in HS skin, cases were stained with multiplexed CD3 and PAX5 IHC to identify T cells and B cells, respectively. PAX5 does not identify terminally differentiated B lineage plasma cells that also accumulate in lesional skin^15^. Digital image quantification of cells on CD3/PAX5 IHC stained tissue sections (Fig. S1) revealed that the frequency of T cells (∼80%) was greater than that of B cells (∼20%) in HS skin but did not differ significantly based on the presence or absence of LA or TLS (Fig. 1D). The accumulation of B cells in HS tissue contrasted with that of normal skin, which only showed an accumulation of T cells with less cellularity than that seen in HS skin. Across all cases, T cells were numerically more plentiful than B cells in HS skin (Fig. 1E, F). Interestingly, the number of T cells and B cells in HS skin trended toward being most sizeable in case with no LA/TLS and lowest in cases with TLS (Fig. 1F). Assessment of CD3/PAX5 IHC in discreet annotated regions of LA and TLS showed a significant skewing toward an increased frequency of B cells in the latter (Fig. 1G). Whereas LA tended to accumulate in the dermis, inversely, TLS tended to arise in the subcutaneous adipose tissue or areas of fibrosis in the subcutis (Fig. 1H). Collectively, these results demonstrate the presence of *bona fide* TLS in HS skin and show that while T cells are the predominant lymphocyte subtype, TLS demonstrate a preponderance of B cells, thereby supporting the hypothesis that B cells may play an important role in the pathogenesis of a subset of patients with this disease.

### B cells in HS skin exhibit signatures of active maturation

To investigate B lymphocyte cell states in HS lesions, we performed scRNA-seq on CD3^-^CD45^+^ cells isolated from five HS patients and five healthy skin donors. B cells were subsetted by *MS4A1* expression and their identity further verified based on differential presence of *CD19, BANK1, CD3E, GNLY, CD14, FCER1A, FCGR3A* (Fig. S2A)^16^. Analysis of our resulting dataset divided B cells into eight separate clusters (Fig. 2A). As the number of B cells derived from healthy skin was comparatively low, we removed these cells from further analysis and focused on identifying populations present in HS lesions. In line with our purification strategy targeting CD45^+^ cells, few fully differentiated plasma cells were present in this dataset (Fig. S2B)^17^. In contrast, a large proportion of sequenced cells were phenotypically characteristic of naive B cells. Cluster 0 was highly enriched in cells expressing *IGHD, CD200*, and *TCL1A* which identify naive cells. (Fig. 2B)^16^. The remaining clusters were associated with activated and memory B cell populations given expression of *CD27, SAMSN1, TNFRSF13B*, and *AIM2* (Fig. 2C)^16,18^. Moreover, naive cells were strongly associated with expression of *IGHM* while memory clusters exhibited preferential isotype class switching to IgA (Fig. 2D).

**Fig 2.**
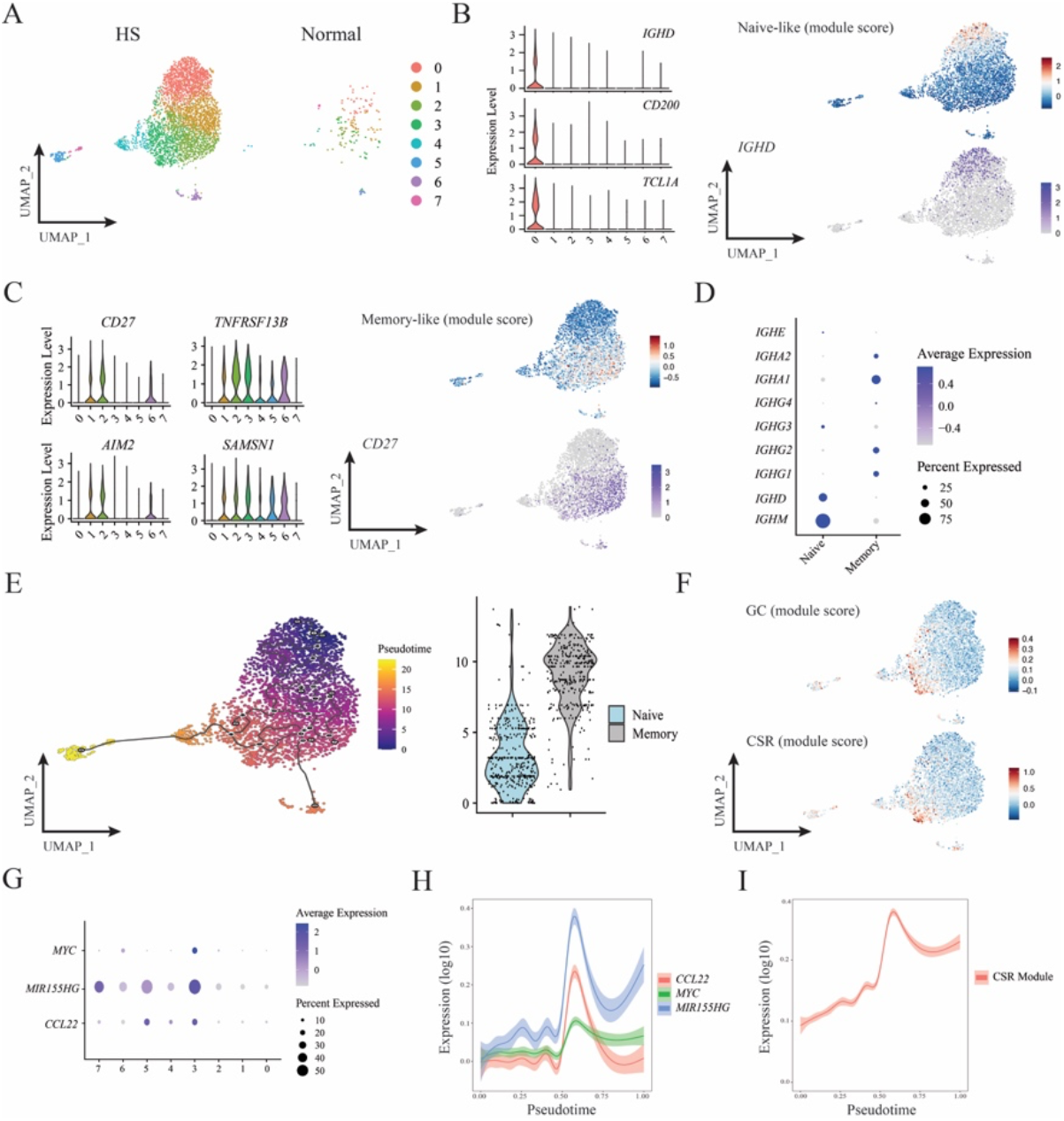
B cells in HS skin exhibit signatures of active maturation. (A) UMAP projections of B cell scRNAseq clusters from either HS lesions or normal normal skin. Data represents concatenation of samples from five HS patents and two normal donors. (B) Expression of naive B cell associated genes and corresponding module score comprising *IGHD, CD200*, and *TCL1A*. UMAP projections depict naive-like module score intensity and *IGHD* expression. (C) Expression of memory B cell associated genes and corresponding module score comprising *CD27, SAMSN1, TNFRSF13B*, and *AIM2*. UMAP projections depict memory-like module score intensity and *CD27* expression. (D) Mean expression of BCR isotype genes among naive and memory B cells from HS lesions. (E) UMAP projection and naive versus memory comparison of B cells from HS lesions analyzed with Monocle 3 for pseudotime. Black lines indicate pseudotime trajectories. (F) UMAP visualization of GC and CSR module scores among B cells from HS lesions. (G) Mean expression of GC positive selection associated genes across HS lesional B cell clusters. (H and I) Plots display gene expression by pseudotime.

Analysis of patient matched blood samples revealed a broadly similar distribution of B cells (Fig. S3A). Both naive and memory populations were equivalently present in HS and normal blood samples and these cells exhibited similar patterns of isotype switching (Fig. S3B, C). Nonetheless, differential expression patterns of several genes implicated in B cell function were evident between HS and healthy blood cells. HS derived blood B cells maintained lower levels of *CD83* and *CD69* while expressing elevated *CD180* and *CCR7* (Fig. S3D). The association of these genes with activation (*CD83, CD69, CD180*)^19–22^ and trafficking (*CCR7*)^23^ is suggestive of systemic qualitative alterations in the B cell compartment of HS patients and consistent with B cell reorganization during inflammation^24^.

Pseudotemporal ordering supported our stratification of cells based on *CD27, CD200*, and *TCL1A* expression. These trajectories arranged memory-like populations progressively later in the trajectories and indicated dynamic relationships between identified clusters (Fig. 2E). Consistent with the presence of TLS identified in our histological analysis, one of the B cell clusters (cluster 3) with intermediary pseudotime values was enriched for genes closely associated with germinal center (GC) B cells^25^. In addition, this cluster exhibited preferential expression of class switch recombination (CSR) machinery genes, including *AICDA, APEX1*, and *XRCC5* (Fig. 2F)^26^. Cluster 3 B cells exhibited hallmarks of active germinal center positive selection. These cells expressed high levels of *MYC, MIR155HG*, and *CCL22* which are important contributors to B cell affinity maturation^27–30^. *CCL22* expression, in particular, overlapped closely with cluster 3 and has been previously defined as a marker of high–affinity germinal center B cells (Fig. 2G)^28^. Across pseuodotime, expression of both these GC positive selection genes (Fig. 2H) and our CSR gene module score (Fig. 2I) peaked at an intermediary value consistent with progressive maturation of naïve cells through the GC towards terminal differentiation as plasma cells. These data demonstrate that a heterogeneous mixture of B cells spanning a spectrum of activation and differentiation cell states accumulate in HS lesions. Altogether, this is highly suggestive of ongoing affinity maturation within lesional skin.

### Spatial transcriptomics identifies distinct regions of TLS involvement in HS skin

In an attempt to definitively characterize B cell organization and potential TLS structures in HS, we analyzed two skin sections from HS lesions and one section of normal skin with spatial transcriptomics (Fig. 3A). Histological analysis revealed pathological changes characteristic of HS, including deep extension of epidermal tracts, widespread immune infiltration, and a histologically identified TLS in the second HS sample (HS2). While HS dermal and epidermal tissue regions clustered transcriptionally with normal skin, regions deeper within HS lesions showed greater organizational disruption and alteration in transcriptional programming (Fig. 3A). Immune cell infiltration, including B cells, was extensive in both HS1 and HS2. B cell lineage markers, *MZB1* and *MS4A1* were expressed in both HS tissues and not detected in normal skin (Fig. 3B). Analysis with published signatures for TLS^31^ and germinal center^25^ activity showed spatially distinct areas bearing these B cell organization signatures in both HS samples, with highest activity occurring within the histologically identified TLS (Fig. 3C). Ligand-receptor interactions within the region of highest TLS/GC activity score of each HS tissue were identified with CellphoneDB (Fig. 3D)^32^. Many HS1 interactions involved MHC Class II genes and chemokines and were thus indicative of antigen presentation and immune cell recruitment. Antigen presentation interactions were also present within HS2; however, interactions driving B cell recruitment (CXCL13_CXCR5)^14^ and germinal center function *via* CD22 engagement^33^ were amongst the top interactions detected. While *CXCL13* expression was more widespread throughout both HS tissues, expression of *CR2*, a highly specific gene for germinal center organizing follicular dendritic cells^34^, was limited to the HS2 TLS cluster (Fig. 3E). Therefore, while discrete TLS regions are readily identified in HS tissues, B cells are more broadly localized within the inflamed skin, and engagement with antigen-presenting cells may occur within and outside of TLS.

**Fig 3.**
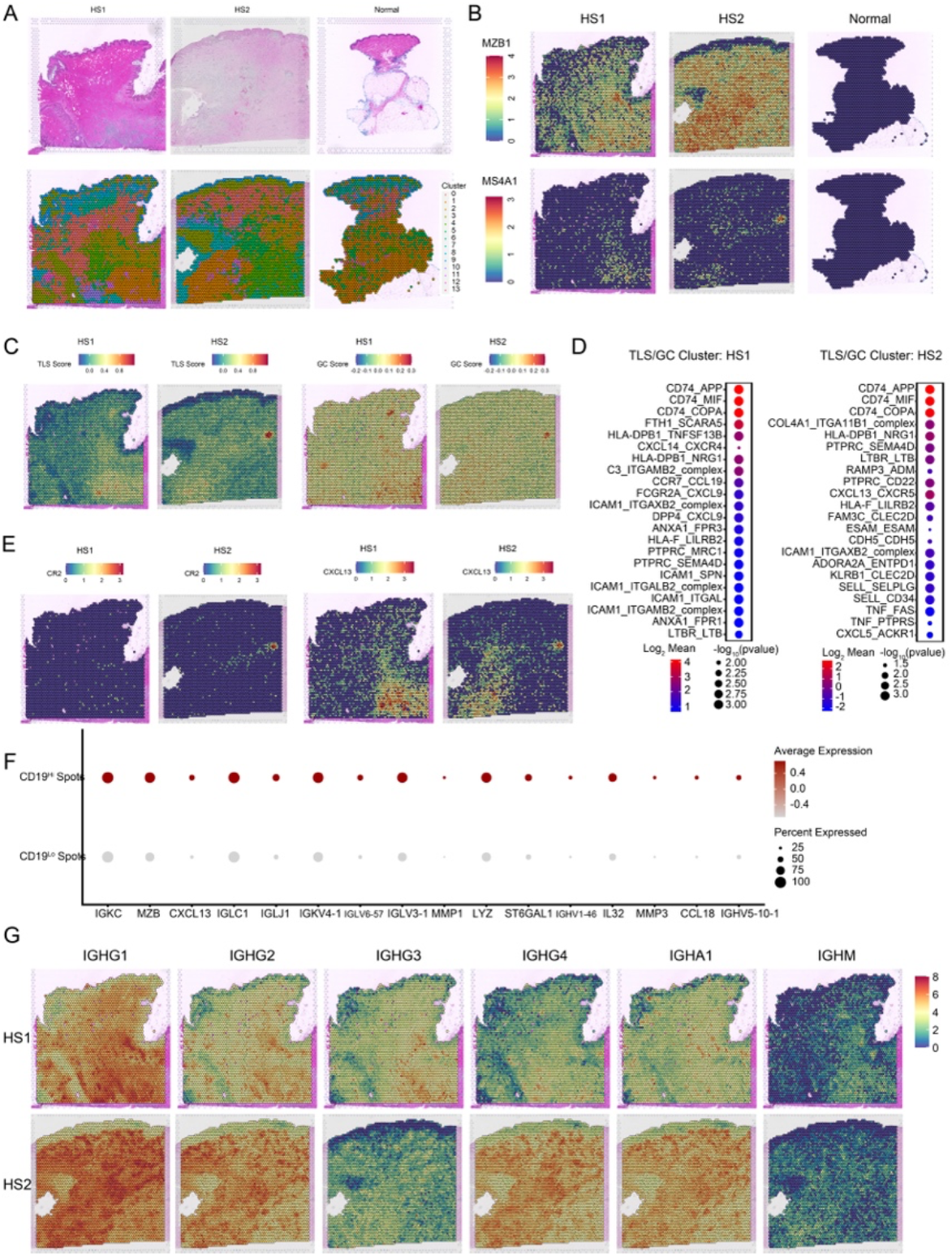
Spatial transcriptomics identifies distinct regions of TLS involvement in HS skin. (A) H&E staining (top) and unsupervised clustering (bottom) of spatial transcriptomics data of two HS samples and one healthy skin sample. (B) Spatial feature plot depicting expression of B cell lineage marker genes *MZB1* and *MS4A1* in spatial transcriptomics data of two HS samples and one healthy skin samples. (C) Spatial feature plot depicting module scores of TLS signature genes (left) and GC signature genes (right) in two HS skin samples. (D) Dot plot of CellphoneDB analysis depicting the top significant ligand: receptor interactions occurring within clusters of high GC/TLS scores in HS samples (cluster 14, HS1, cluster 8, HS2) (p<0.05). (E) Spatial feature plot depicting expression of B cell recruitment factor *CXCL13* and follicular dendritic cell marker *CR2* in two HS skin samples. (F) Dot plot depicting significantly increased genes comparing spots with high expression of *CD19* (*CD19* >= 0.5) versus low expression of CD19 (*CD19* < 0.5) (adjusted p<0.05). (G) Spatial feature plot depicting expression of immunoglobulin genes *IGHG1, IGHG2, IGHG3, IGHG4, IGHA1*, and *IGHM* in two HS skin samples.

To understand how B cells might be affecting skin tissue, we asked what transcriptional differences were increased in tissue spots containing high levels of *CD19* transcript versus low transcript signals (Fig. 3F). Most differentially expressed genes were in the immunoglobulin family, with B lineage marker *MZB1* and B cell recruitment factor *CXCL13* also differentially expressed. Interestingly, several genes associated with wound repair, including *MMP1, MMP3*, and *IL32* were also enriched in B cell containing spots^35,36^. Given that immunoglobulin genes are differentially expressed within B cell containing spots, we assessed expression of specific B cell isotypes within the tissues (Fig. 3G). IgG1 expression was prevalent amongst both tissues, while IgG2, IgG3, and IgG4 expression varied amongst individual patients. IgA, a key mediator of mucosal tissue protection, was also highly detected in both tissues^37^. In contrast, IgM expression was lowest in both tissues, indicating that the majority of antibody producing B lineage cells had class-switched. Thus, within HS lesional tissue, immunoglobulin production, potentially around areas with heightened wound repair signals, is a predominant cellular process in B cells and plasma cells.

### Fibroblasts and T cells are primed to recruit and support B cells in HS skin

Next, we asked whether tissue stromal cells play a role in recruiting and supporting B cell activity in HS skin. We performed scRNASeq on purified CD45 negative cells from three HS lesional skin specimens and six normal skin specimens, which clustered into subsets composed of fibroblasts (marked by *PDPN, DCN*, and *PDGFRA*) and endothelial cells (marked by *PECAM1, CDH5*, and *TIE1*) (Fig. 4A). Given the potential role of keratinocytes in HS pathology, we additionally sequenced CD45 negative cells from epidermal preparations of two HS specimens and two healthy skin specimens (Fig. 4B). CellphoneDB analysis between HS skin endothelial cell clusters, fibroblast clusters, keratinocyte clusters and B cells identified numerous interactions between stromal cell receptor/ligands and B cells (Fig. 4C). Of note, CXCL13 engagement of CXCR5 was solely identified in fibroblast clusters. Since candidate cell interactions were widespread across numerous cell types, we then asked which stromal cell derived interaction partners were differentially expressed between HS and normal skin cell types. While most interaction partners expressed by endothelial cells were also expressed in healthy skin (Supplemental Fig. S4A), fibroblasts in HS skin upregulated numerous molecules versus healthy controls (Fig. 4D).Amongst these were chemokines *CXCL13, CXCL10, CCL3*, and prostaglandin receptor family members. Expression of these factors were not confined to a single fibroblast cluster and were lower or absent from normal skin (Fig. 4E). While keratinocytes also upregulated some chemokines in HS skin, these were limited to lower abundance subsets (Supplemental Fig. S4B). Therefore, fibroblasts may play a key role in recruiting and supporting B cells in HS tissues.

**Fig 4.**
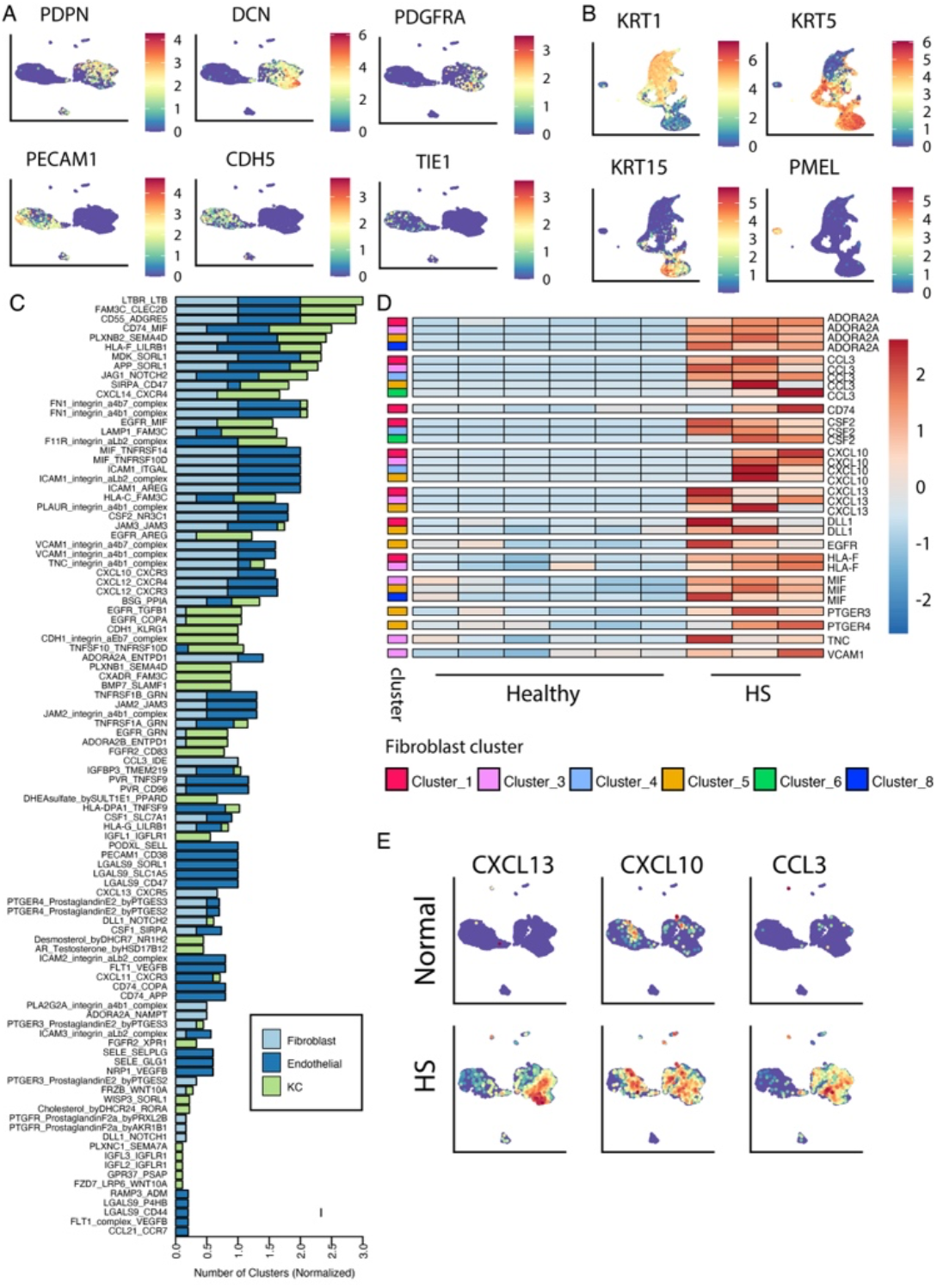
Fibroblasts are primed to support and recruit B cells in HS skin. (A) UMAP projections of stromal cell scRNASeq clusters from HS lesions and normal skin showing expression of endothelial and fibroblast signature genes. Data represents samples from three HS donors and 6 normal skin donors. (B) UMAP projections of scRNASeq data of epidermal cells from HS lesions and normal skin showing expression of keratinocyte signature genes. Data represents samples from two HS donors and 2 normal skin donors. (C) Significant (p<0.005) ligand:receptor interaction partners identified by CellphoneDB between clusters of stromal cells (left partner) and B cells (right partner). Numbers of clusters involved are normalized to the total number of clusters involved per cell type. (D) Row-normalized heatmap depicting pseudo-bulk scRNASeq counts of significantly upregulated (adjusted p<0.05) genes in HS fibroblast clusters versus healthy skin fibroblast clusters. (E) UMAP projections of stromal cell scRNASeq clusters from HS lesions and normal skin showing expression of select differentially expressed genes. Data represents samples from three HS donors and 6 normal skin donors.

CD4^+^ T peripheral helper cells (Tph) are critical for the formation of functional TLS^14,38^. These cells closely resemble the T follicular helper cells populating lymphoid organs and promote B cell affinity maturation through provision of co-stimulation and supportive cytokines^38^. To determine if Tph were present in HS lesions we analyzed CD4^+^ T cell populations with scRNAseq (Fig. 5A). CellphoneDB analysis between CD4^+^ T cell clusters and B cells identified numerous interactions across clusters, especially *CXCL13* and the B cell survival factor *TNSF13B* (BAFF) (Fig. 5B). We next generated a “Tph/Tfh” signature score incorporating the genes most highly expressed by these cells^39^. Tph genes were highly expressed by CD4^+^ T cell cluster 6 and this population was enriched in HS samples compared to healthy controls (Fig. 5C). In addition, “Germinal Center T Helper Up” was the among the top 10 pathways significantly defined by Gene Set Enrichment Analysis (GSEA) comparing cluster 6 to all other T cells (Fig. 5D)^40^. Analysis of the clusters with the strongest Tph signature (clusters 3, 6, 10) revealed increased expression of canonical Tph genes, especially *CXCL13, TOX2*^41^, *TCF7*^42^, and *KLRB1*^43^, compared to healthy controls.

**Fig. 5.**
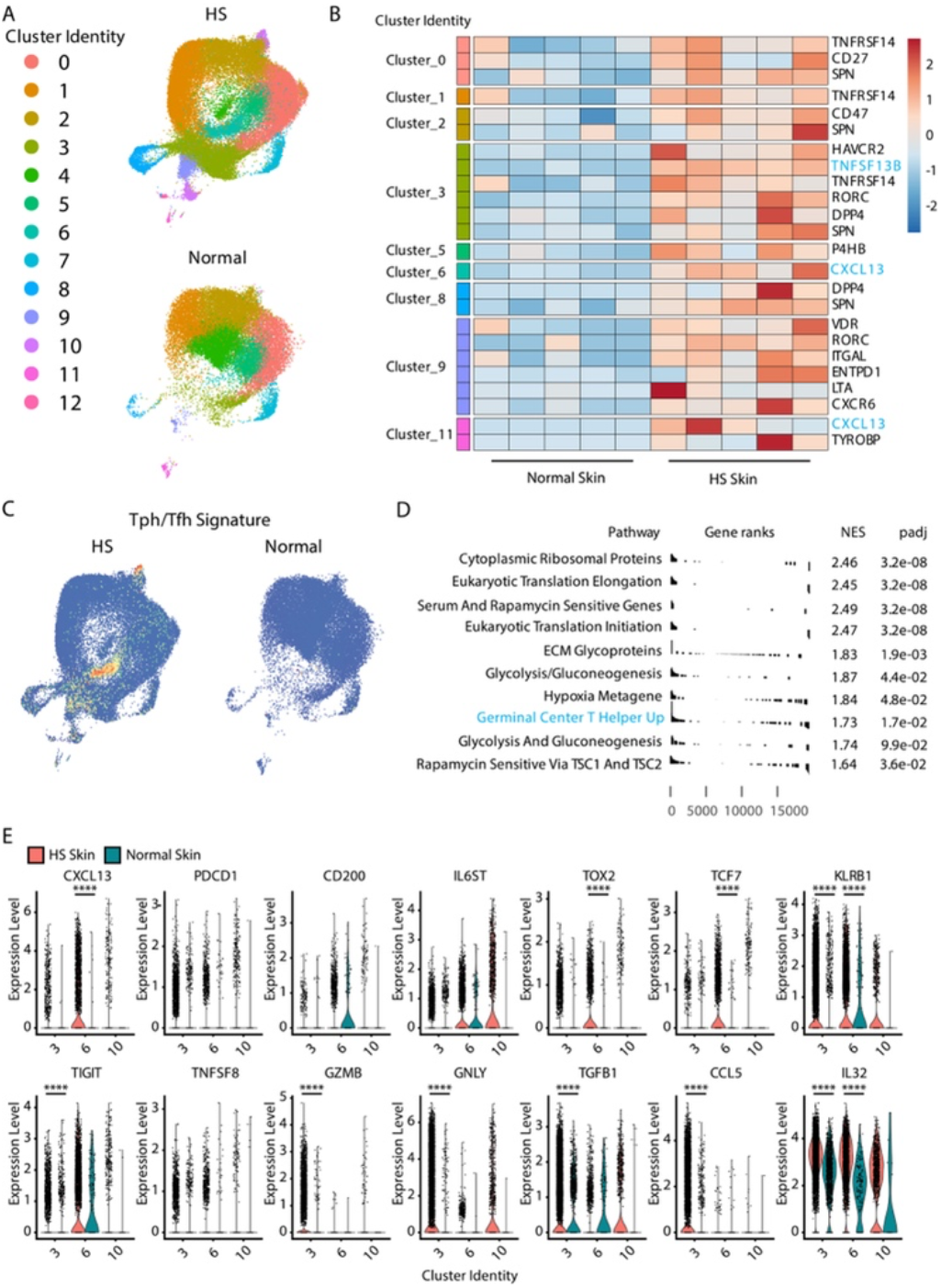
CXCL13-expressing T peripheral helper cells are increased in HS lesional skin. (A) UMAP projections of CD4 T cell scRNASeq clusters from HS lesions and normal skin. (B) Row-normalized heatmap depicting pseudo-bulk scRNASeq counts of significantly upregulated (adjusted p<0.05) genes identified as B cell interaction partners in HS CD4 T cell clusters versus healthy skin CD4 T cell clusters. (C) UMAP projections of CD4 T cell scRNASeq clusters from HS lesions and normal skin showing expression of a Tph/Tfh gene module. (D) Top ten significantly enriched (padj<0.05) GSEA results of Human MSigDB Collections C2 gene signatures comparing cluster 6 cells from HS lesional skin to all other clusters. (E) Violin plots of gene expression from clusters 3, 6, and 10 from HS lesions and normal skin showing expression of select genes. Data represents samples from five HS donors and five normal skin donors. **** p<0.0001

Cluster 3 was also enriched in the cytotoxic genes *GZMB* and *GNLY* which have been associated with Tph/Tfh regulation (Fig. 5E)^44^. The cytokine *IL32*, which has been implicated as a potential plasma cell survival factor, was enriched in both clusters 3 and 6 (Fig. 5E) as well as significantly elevated in CD19^hi^ spots (Fig. 3F). Altogether, our analyses of the T cell and stromal compartment in HS lesions indicates that specific populations of these cells, including Tph, are expanded and actively express genes associated with TLS function and B cell maturation.

## Discussion

While B cell signatures are strongly evident in HS, the contribution of these lymphocytes to disease pathogenesis is unclear. Memory B cells and antibody secreting plasma cells accumulate in HS lesional skin to become a prominent immunological feature^1,4–6^. Antibodies against a wide array of self-antigens are detectable in both patient sera and the skin lesions themselves suggesting ongoing B cell activity and possible contribution to disease progression^6–9^. Given healthy skin maintains relatively few B cells it is not clear how HS skin lesions supports accumulation of these cells and whether specific tissue niches regulate their biological activity. Our data demonstrate the presence of TLS, capable of supporting and shaping B cell function, within HS lesions. We find evidence of active B cell maturation and affinity maturation within lesional skin and identify a cellular niche of skin stromal constituents capable of B cell recruitment and maintenance.

TLS are organized aggregates of immune cells that form in nonlymphoid tissues, especially under chronic inflammatory conditions. TLS are composed of T cells, B cells, myeloid antigen presenting cells and supporting stromal cells coalescing to resemble secondary lymphoid organs. As such, these structures are capable of regulating B cell immune responses by facilitating T cell help and supporting formation of germinal center reactions^14^. Lymphoid aggregates are recognized as a histological feature of HS lesions and the presence of fully formed TLS has been hypothesized; however, definitive evidence of fully functional TLS has not previously been reported^1,4,13^. Our data demonstrate that these structures are relatively common in advanced HS lesions. Moreover, within these TLS we identified evidence of active germinal centers and B cells expressing genes associated with affinity maturation and class switch recombination. A significant portion of detected cells were class switched to IgA which is associated with mucosal and epithelial surfaces^37^. We observed that B cells accumulating in HS lesions contain significant population heterogeneity with evident recruitment of both naive and memory cells. Due to the limitations of our sample cohort, we were unable to quantify TLS frequency or identify correlations with unique patient subsets. Nonetheless, these data point to a model where B cell activity is propagated within the cutaneous tissue and accumulating lesional B cells likely mature to generate fully differentiated plasma cells *in situ*. This would be consistent with the relationship between TLS B cells and tissue plasma cells described in other situations of chronic inflammation. In renal cell carcinoma tumor TLS generate IgA and IgG plasma cells that disseminate relatively large distances across the tumor tissue^45^. While we have not uncovered a direct link between HS TLS B cells and production of autoantibodies, it is plausible that cutaneous germinal centers give rise to the plasma cells secreting these antibodies.

Naive B cells are rare in healthy skin and therefore likely recruited to progressing HS lesions through the induction of a supportive tissue microenvironment complete with associated chemokines and maintenance factors^46^. In agreement with this model, we and others have previously observed significant upregulation of the B cell survival factors BAFF and APRIL in HS lesions^4,5^. That organized TLS are also induced by the HS inflammatory milieu provides further evidence for generation of a niche supportive of B cells during the progressive evolution of HS lesions. Our data demonstrate that several chemokines associated with B cell recruitment to TLS, especially CXCL13, CCL10, and CCL18, are enriched in lesional skin^14,31,46,47^. Notably, fibroblasts were the only stromal cell population that expressed *CXCL13* despite the high levels of transcript detected. This indicates a key role for fibroblasts in B cell recruitment and propagation of TLS activity.

To date, limited data suggest that targeting B cells may be beneficial in the treatment of HS. B cell depletion with rituximab ameliorated HS lesions in a single patient and reduced expression of inflammatory cytokines in HS skin explant assays^11,12^. Anti-TNF? agents are the most effective targeted biologics currently available for HS treatment^1^. Notably, TNF? is a key contributor in TLS induction and we have previously demonstrated that *ex vivo* anti-TNF? treatment of lesional skin samples reduced B cell proliferation^5,14,46^. Herein, we present findings that add to the growing evidence suggesting a role for B cells in HS pathogenesis and that immunotherapy targeting these cells may be beneficial for HS patients. Our data imply that plasma cell depletion may be insufficient because cutaneous TLS will be capable of replacing cells secreting pathogenic antibodies. Instead, direct manipulation of TLS formation in HS may allow for attenuation of pathogenic B cell activity while reducing systemic impacts on healthy tissues.

## Methods

### Study approval

The UCSF Institutional Review Board approved the proposed studies (13-11307, 19-29608, and 21-33678). All patients provided informed written consent. Keratinocyte cells were isolated from discarded, de-identified tissue certified as Not Human Subjects Research.

### Case Selection

Thirty cases of HS were retrieved in the diagnostic and consultation files of the UCSF Dermatopathology Service and UCSF Department of Surgical Pathology. Of these, 3 cases of HS with germinal center reactions, 3 cases with lymphoid aggregates but no germinal center reaction, and 3 cases of no germinal center reactions or lymphoid aggregates were selected for immunohistochemical studies. Notably, one case initially selected as having only lymphoid aggregates and no germinal center reactions on hematoxylin and eosin (H&E) staining exhibited a FDC network on CD21 IHC and was therefore re-classified as an HS case with a TLS. One case of microscopically near normal skin from an anogenital site was selected from UCSF Dermatopathology Service.

### Histology

Tissue was fixed in 10% neutral-buffered formalin, routinely processed, embedded in paraffin, and stained with H&E. Formalin-fixed paraffin-embedded (FFPE) sections of 4 μm thickness were stained with antibodies specific for CD21 (predilute, clone 2G9, Leica Biosystems, Danvers, MA), CD3 (predilute, clone LN10, Leica Biosystems, Danvers, MA), and PAX5 (predilute, clone DAK-Pax5, Agilent Technologies, Santa Clara, CA).

### Whole Slide Digital Image Analysis

Slides were scanned at ×40 resolution with an Aperio AT2 scanner (Leica Biosystems Danvers, MA) using a 20×/0.75NA Plan Apo objective with a 2× optical magnification changer. Quantitative analysis was performed on images from sections stained with a dual antibody combination of PAX5 (red chromogen) and CD3 (brown chromogen) using QuPath (v.0.3.2) software. Small representative regions of interest of hematoxylin, 3,3’-diaminobenzidine (CD3), alkaline phosphatase (PAX5) and background were used to set stain vectors. Cells were detected using the watershed algorithm-based Detect Cells tool, using the optical density sum to segment nuclei with a cytoplasmic expansion of 1.5 µm. Cells were then classified with the Train Object Classifier tool using a neural network trained on 8–12 annotations of each cell class (CD3^+^PAX5^-^, CD3^-^PAX5^+^, and CD3^-^PAX5^-^) with the following features as input: nuclear PAX5 mean and cytoplasmic CD3 mean. LA and TLS were manually annotated for cell localization, and tissue area was detected with a simple thresholder.

### Spatial Transcriptomics

Formalin fixed paraffin embedded specimens from two lesional HS samples and one healthy control sample were sectioned and placed on a Visium Spatial Gene Expression Slide (10x Genomics). Probesets were amplified and were sequenced by the Gladstone institute. Fastq files and the haematoxylin and eosin stained image were analyzed by spaceranger (v. 1.3.1) using the Visium Human Transcriptome Probe Set v1.0 (GRCh38 2020 A). Spaceranger output files were analyzed with Seurat^48^. Data was normalized with SCTransform prior to PCA and UMAP clustering^49^. For analysis comparing multiple slides, separate slides were merged and integrated with Harmony (v. 0.1.0)^50^ prior to clustering. GC and TLS scores were calculated using the AddModuleScore function using gene sets described for germinal centers^25^ and tertiary lymphoid structures^31^ (supplemental file: SupplementaryTableGeneLists.xlsx).

Spots with high expression of CD19 (CD19>= 0.5) were compared to spots with low CD19 expression (CD19<0.5) using the FindMarkers command, and significantly increased genes (p_val_adj <0.05) were plotted.

Counts assigned to clusters were exported from each HS object and analyzed with cellphoneDB^32^. The top significant interactions (p<0.05) occurring within clusters of local high GC/TLS scores (cluster 14, HS1 (mean>1), Cluster 8, HS2 (mean>0) were identified and plotted.

### Skin Processing

6mm punch biopsies were obtained from five patients with a diagnosis of hidradenitis suppurativa from an actively inflamed lesion. Healthy control skin was obtained from surgical discards and was dermatomed at 500 micron prior to processing. Skin was finely minced with scissors and was digested overnight with 250 IU/mL collagenase IV (Worthington) and DNAse 20 µg/mL (Sigma) in RPMI with 10 %FBS at 37 deg C. The suspension was agitated via shaking and was filtered (100 μm) and washed before counting with a NucleoCounter NC-200 (ChemoMetec). Samples were stained for sort-purification with an Aria2 with Ghost Dye Violet 510 amine reactive dye (Tonbo), CD45 PerCP-e710, CD3 BV650, CD4 PE-CF594, CD127 PE, CD25 PE-Cy7, CD45RO FITC, CD27 APC-e780, and CD8 APC. Some samples were also stained with CD235a Pacific Blue.

CD45 negative cells were sorted as singlet, Ghost Dye negative, CD45 negative events. Some samples were also sorted as CD235a negative to exclude erythrocytes. CD3 negative events were sorted as singlet, Ghost Dye negative, CD45 positive, CD3 negative events. Regulatory T cells were enriched from skin by sorting as singlet, Ghost Dye negative, CD45 positive, CD3 positive, CD8 negative, CD4 positive, CD25 positive and CD27 positive. CD4 T cells not falling in the CD25 positive and CD27 positive regulatory T cell gate were also collected. Regulatory T cells and non-regulatory T cells were either spiked in a 1:1 ratio and sequenced together or sequenced on separate 10x wells.

### Keratinocyte preparation

Discarded skin from two HS surgical excisions and two healthy skin resections was dermatomed at 1000 micron depth and suspended overnight in a solution of 5U/mL dispase (Stemcell cat #07913) at 4deg C. The epidermis was separated mechanically the next day, and was placed in a 0.25% trypsin-EDTA solution for 15 minutes at 37 deg C. The epidermis was chopped with scissors and passed through a 100 µ filter, washed, and stained with anti-CD45-PerCP710, anti-CD3-BV711, anti-CD19-FITC, and Ghost Dye Violet 510 prior to sort-purification with an Aria Fusion. Keratinocytes were enriched for as singlet, live, CD45 negative events.

### Single Cell RNA sequencing

Following sort purification, CD45 negative and CD3 negative cells were quantified via nucleocounter or hemacytometer and 10,000 to 25,000 cells were loaded per lane and sequenced with a 10x Genomics Single Cell 5′ chip. Keratinocytes (25,000) were loaded onto a 10x Genomics Single Cell 3′ chip. All sample loading and sequencing was performed by the UCSF Genomics Core Facility, which provided fastq files following sequencing.

Fastq files were processed via cellranger (v. 3.0.2) to transcriptome GRCh38 v 3.0.0 downloaded from 10x Genomics.

### Seurat analysis

Cellranger generated counts files were analyzed with Seurat (4.1.0)^48^. First, singlets were identified with scDblFinder (1.4.0)^51^. Singlets were further selected as cells containing >500 nFeature_RNA and less than 5% mitochondrial reads. Harmony (v. 0.1.0)^50^ was used for sample integration and clustering, with CD3 negative sorted cells, CD45 negative sorted cells, and keratinocytes and T cells integrated into four separate objects for analysis.

B cell clusters were identified from the CD3 negative Seurat object based upon *MS4A1* expression (threshold) and were subset and reclustered. B cell purity was confirmed based on the differential presence of *CD19, BANK1, CD3E, GNLY, CD14, FCER1A, FCGR3A*. Trajectory and pseudotime analysis of B cell populations was performed with Monocle 3^52^. GC and CSR scores were calculated using the AddModuleScore function and gene sets described for germinal centers^25^ and active class switch recombination^26^ (supplementary file: SupplementaryTableGeneLists.xlsx).

### CellphoneDB analysis

CellphoneDB (v. 4.0.0)^32^ was used to identify ligand-receptor interactions between fibroblasts, endothelial, keratinocytes, and B cells in HS tissues or between CD4 T cells and B cells in HS tissues. Normalized counts from HS clusters were exported and assessed with the cellphoneDB statistical_analysis method. Any interaction between a B cell cluster and a keratinocyte/endothelial/fibroblast cluster or between a B cell cluster and a CD4 T cell that was significant (p<0.005) was identified. For stromal cell types, numbers of interactions between cell types were counted, normalizing for the total number of clusters identified per cell type. Differential expression analysis was performed with DESeq2 (v. 1.30.1)^53^ on pseudobulk counts for individual patient clusters, comparing cells from HS patients to healthy controls. Genes that were significantly increased in HS clusters versus controls were depicted with pheatmap (v. 1.0.12).

### Gene Set Enrichment Analysis

A ranked gene list for GSEA was generated to compare expression between cluster 6 and all other HS T cell clusters. First, all HS CD4 T cells were downsampled to 10,000 events. The Seurat FindMarkers command was used to determine differential expression between cluster 6 and all other clusters. To obtain rankings for all genes, the logfc.threshold and min.pct were set to 0. fgsea (v. 1.16.0) was used to determine enrichment on the MSigDB C2 “Curated Gene Sets” Collection. Significant (padj<0.05) positively enriched pathways containing more than thirty genes were selected and collapsed into independent pathways with the collapsePathways command. GSEA Tables for the top 10 enriched pathways were plotted.

## References

1. van Straalen, K. R., Prens, E. P. & Gudjonsson, J. E. Insights into hidradenitis suppurativa. J. Allergy Clin. Immunol. 149, 1150–1161 (2022).

2. Yu, C. C. & Cook, M. G. Hidradenitis suppurativa: a disease of follicular epithelium, rather than apocrine glands. Br. J. Dermatol. 122, 763–769 (1990).

3. Navrazhina, K. et al. Epithelialized tunnels are a source of inflammation in hidradenitis suppurativa. J. Allergy Clin. Immunol. 147, 2213–2224 (2021).

4. Sabat, R. et al. Neutrophilic granulocyte-derived BAFF supports B cells in skin lesions in hidradenitis suppurativa. J. Allergy Clin. Immunol. (2022).

5. Lowe, M. M. et al. Immunopathogenesis of hidradenitis suppurativa and response to anti-TNF-α therapy. JCI Insight 5, (2020).

6. Gudjonsson, J. E. et al. Contribution of plasma cells and B cells to hidradenitis suppurativa pathogenesis. JCI Insight 5, (2020).

7. Assan, F. et al. Anti-Saccharomyces cerevisiae IgG and IgA antibodies are associated with systemic inflammation and advanced disease in hidradenitis suppurativa. J. Allergy Clin. Immunol. 146, 452-455.e5 (2020).

8. Carmona-Rivera, C. et al. Autoantibodies Present in Hidradenitis Suppurativa Correlate with Disease Severity and Promote the Release of Proinflammatory Cytokines in Macrophages. J. Invest. Dermatol. 142, 924–935 (2022).

9. Hoffman, L. K. et al. Integrating the skin and blood transcriptomes and serum proteome in hidradenitis suppurativa reveals complement dysregulation and a plasma cell signature. PLoS One 13, e0203672 (2018).

10. Macchiarella, G. et al. Disease Association of Anti-Carboxyethyl Lysine Autoantibodies in Hidradenitis Suppurativa. J. Invest. Dermatol. (2022).

11. Vossen, A. R. J. V., Ardon, C. B., van der Zee, H. H., Lubberts, E. & Prens, E. P. The anti-inflammatory potency of biologics targeting tumour necrosis factor-α, interleukin (IL)-17A, IL-12/23 and CD20 in hidradenitis suppurativa: an ex vivo study. Br. J. Dermatol. 181, 314–323 (2019).

12. Takahashi, K. et al. Successful treatment of hidradenitis suppurativa with rituximab for a patient with idiopathic carpotarsal osteolysis and chronic active antibody-mediated rejection. J. Dermatol. 45, e116–e117 (2018).

13. van der Zee, H. H. et al. Alterations in leucocyte subsets and histomorphology in normal-appearing perilesional skin and early and chronic hidradenitis suppurativa lesions. Br. J. Dermatol. 166, 98–106 (2012).

14. Schumacher, T. N. & Thommen, D. S. Tertiary lymphoid structures in cancer. Science 375, eabf9419 (2022).

15. Nera, K.-P. et al. Loss of Pax5 promotes plasma cell differentiation. Immunity 24, 283–293 (2006).

16. Stewart, A. et al. Single-Cell Transcriptomic Analyses Define Distinct Peripheral B Cell Subsets and Discrete Development Pathways. Front. Immunol. 12, 602539 (2021).

17. Caraux, A. et al. Circulating human B and plasma cells. Age-associated changes in counts and detailed characterization of circulating normal CD138- and CD138 plasma cells. Haematologica vol. 95 1016–1020 Preprint at https://doi.org/10.3324/haematol.2009.018689 (2010).

18. Weisel, N. M. et al. Surface phenotypes of naive and memory B cells in mouse and human tissues. Nat. Immunol. 23, 135–145 (2022).

19. Prazma, C. M., Yazawa, N., Fujimoto, Y., Fujimoto, M. & Tedder, T. F. CD83 expression is a sensitive marker of activation required for B cell and CD4+ T cell longevity in vivo. J. Immunol. 179, 4550–4562 (2007).

20. Ashouri, J. F. & Weiss, A. Endogenous Nur77 Is a Specific Indicator of Antigen Receptor Signaling in Human T and B Cells. J. Immunol. 198, 657–668 (2017).

21. Chaplin, J. W., Chappell, C. P. & Clark, E. A. Targeting antigens to CD180 rapidly induces antigen-specific IgG, affinity maturation, and immunological memory. J. Exp. Med. 210, 2135–2146 (2013).

22. Chaplin, J. W., Kasahara, S., Clark, E. A. & Ledbetter, J. A. Anti-CD180 (RP105) activates B cells to rapidly produce polyclonal Ig via a T cell and MyD88-independent pathway. J. Immunol. 187, 4199–4209 (2011).

23. Förster, R. et al. CCR7 Coordinates the Primary Immune Response by Establishing Functional Microenvironments in Secondary Lymphoid Organs. Cell vol. 99 23–33 Preprint at https://doi.org/10.1016/s0092-8674(00)80059-8 (1999).

24. Moreau, J. M. et al. Inflammation rapidly reorganizes mouse bone marrow B cells and their environment in conjunction with early IgM responses. Blood 126, 1184–1192 (2015).

25. Victora, G. D. et al. Identification of human germinal center light and dark zone cells and their relationship to human B-cell lymphomas. Blood 120, 2240–2248 (2012).

26. King, H. W. et al. Single-cell analysis of human B cell maturation predicts how antibody class switching shapes selection dynamics. Sci Immunol 6, (2021).

27. Thai, T.-H. et al. Regulation of the germinal center response by microRNA-155. Science 316, 604–608 (2007).

28. Liu, B. et al. Affinity-coupled CCL22 promotes positive selection in germinal centres. Nature 592, 133–137 (2021).

29. Dominguez-Sola, D. et al. The proto-oncogene MYC is required for selection in the germinal center and cyclic reentry. Nat. Immunol. 13, 1083–1091 (2012).

30. Calado, D. P. et al. The cell-cycle regulator c-Myc is essential for the formation and maintenance of germinal centers. Nat. Immunol. 13, 1092–1100 (2012).

31. Wu, R. et al. Comprehensive analysis of spatial architecture in primary liver cancer. Sci Adv 7, eabg3750 (2021).

32. Efremova, M., Vento-Tormo, M., Teichmann, S. A. & Vento-Tormo, R. CellPhoneDB: inferring cell–cell communication from combined expression of multi-subunit ligand– receptor complexes. Nat. Protoc. 15, 1484–1506 (2020).

33. Meyer, S. J. et al. CD22 Controls Germinal Center B Cell Receptor Signaling, Which Influences Plasma Cell and Memory B Cell Output. J. Immunol. 207, 1018–1032 (2021).

34. Kranich, J. & Krautler, N. J. How Follicular Dendritic Cells Shape the B-Cell Antigenome. Front. Immunol. 7, 225 (2016).

35. Caley, M. P., Martins, V. L. C. & O’Toole, E. A. Metalloproteinases and Wound Healing. Adv. Wound Care 4, 225–234 (2015).

36. Nold-Petry, C. A. et al. IL-32 promotes angiogenesis. J. Immunol. 192, 589–602 (2014).

37. Chen, K., Magri, G., Grasset, E. K. & Cerutti, A. Rethinking mucosal antibody responses: IgM, IgG and IgD join IgA. Nat. Rev. Immunol. 20, 427–441 (2020).

38. Yoshitomi, H. Peripheral Helper T Cell Responses in Human Diseases. Front. Immunol. 13, 946786 (2022).

39. Dunlap, G. S. et al. Single-cell transcriptomics reveals distinct effector profiles of infiltrating T cells in lupus skin and kidney. JCI Insight 7, (2022).

40. Subramanian, A. et al. Gene set enrichment analysis: a knowledge-based approach for interpreting genome-wide expression profiles. Proc. Natl. Acad. Sci. U. S. A. 102, 15545–15550 (2005).

41. Xu, W. et al. The Transcription Factor Tox2 Drives T Follicular Helper Cell Development via Regulating Chromatin Accessibility. Immunity 51, 826-839.e5 (2019).

42. Choi, Y. S. et al. LEF-1 and TCF-1 orchestrate T(FH) differentiation by regulating differentiation circuits upstream of the transcriptional repressor Bcl6. Nat. Immunol. 16, 980–990 (2015).

43. Braud, V. M. et al. LLT1-CD161 Interaction in Cancer: Promises and Challenges. Front. Immunol. 13, 847576 (2022).

44. Xie, M. M. et al. Follicular regulatory T cells inhibit the development of granzyme B-expressing follicular helper T cells. JCI Insight 4, (2019).

45. Meylan, M. et al. Tertiary lymphoid structures generate and propagate anti-tumor antibody-producing plasma cells in renal cell cancer. Immunity 55, 527-541.e5 (2022).

46. Asam, S., Nayar, S., Gardner, D. & Barone, F. Stromal cells in tertiary lymphoid structures: Architects of autoimmunity. Immunological Reviews vol. 302 184–195 Preprint at https://doi.org/10.1111/imr.12987 (2021).

47. Liu, Z., Meng, X., Tang, X., Zou, W. & He, Y. Intratumoral tertiary lymphoid structures promote patient survival and immunotherapy response in head neck squamous cell carcinoma. Cancer Immunology, Immunotherapy Preprint at https://doi.org/10.1007/s00262-022-03310-5 (2022).

48. Hao, Y. et al. Integrated analysis of multimodal single-cell data. Cell 184, 3573-3587.e29 (2021).

49. Hafemeister, C. & Satija, R. Normalization and variance stabilization of single-cell RNA-seq data using regularized negative binomial regression. Genome Biol. 20, 296 (2019).

50. Korsunsky, I. et al. Fast, sensitive and accurate integration of single-cell data with Harmony. Nat. Methods 16, 1289–1296 (2019).

51. Germain, P.-L., Lun, A., Garcia Meixide, C., Macnair, W. & Robinson, M. D. Doublet identification in single-cell sequencing data using scDblFinder. F1000Res. 10, 979 (2021).

52. Cao, J. et al. The single-cell transcriptional landscape of mammalian organogenesis. Nature 566, 496–502 (2019).

53. Love, M. I., Huber, W. & Anders, S. Moderated estimation of fold change and dispersion for RNA-seq data with DESeq2. Genome Biol. 15, 550 (2014).

