## Supplemental Figures for "Tertiary Lymphoid Structures Sustain Cutaneous B cell Activity in Hidradenitis Suppurativa"

**Supplement Figures:**

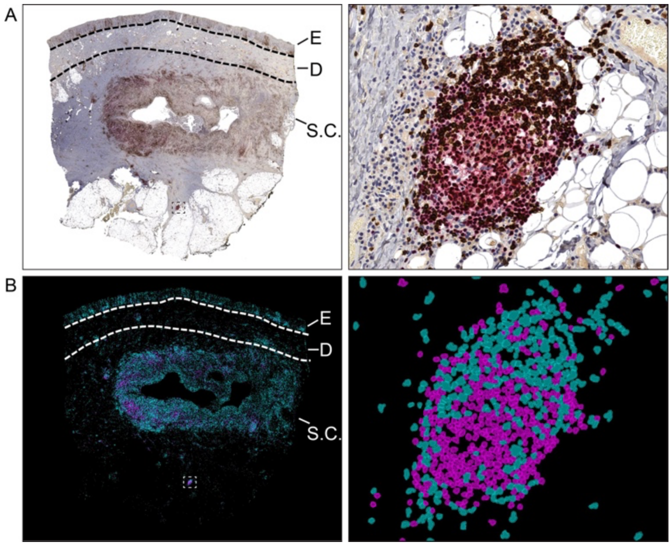


**Fig S1.** Digital image quantification of lymphocytes in skin. (A) Representative low power photomicrograph (left) and high power tertiary lymphoid structure (TLS; right) of the multiplex CD3 (brown chromogen)/PAX5 (red chromogen) immunohistochemical stain on HS skin. (B) Low and high power images corresponding to (A), in which the image analysis program created cell contours according to a cell segmentation algorithm and labeled cells according to signal intensity thresholds set for the red and brown chromogens. Cyan corresponds to CD3+ cells and Magenta corresponds to PAX5+ cells. E: epidermis, D: dermis, S.C.: subcutis. Dashed box indicates the location of TLS highlighted in the high power image.

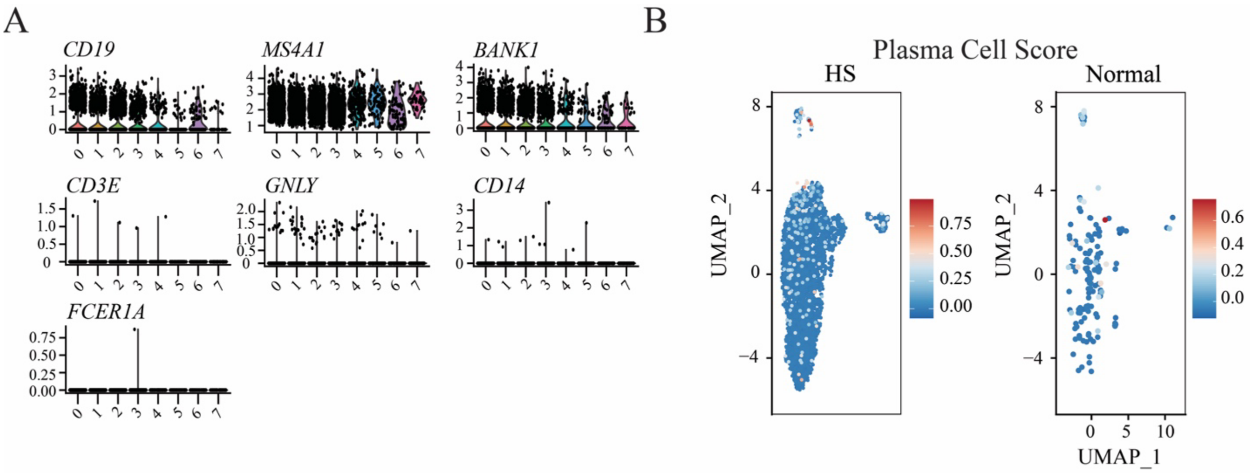


**Fig S2.** Single cell RNA sequencing analysis of B cells from skin. (A) Expression of lineage defining genes among *MS4A1* selected scRNA-seq cell clusters from HS lesions. (B) UMAP visualizations of plasma cell module score (*CD38*, *SDC1*, *TNFRSF17*, *MKI67*, *PRDM1*) intensity in HS lesional and normal skin B cells.

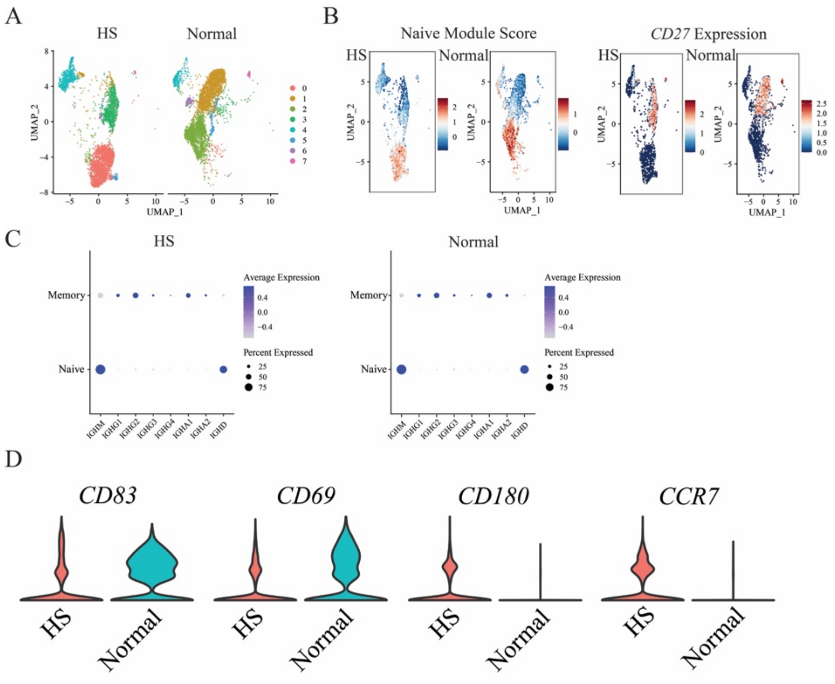


**Fig S3.** Comparative single cell RNA sequencing analysis of healthy and HS blood B cells. (A) UMAP projects of HS and Normal blood B cells. Data represents concatenation of samples from five HS patents and two normal donors. (B) UMAP projections depict naive-like module and *CD27* expression. (C) Mean expression of BCR isotype genes among naive and memory blood B cells from HS or healthy patients. (D) Expression of notable B cell functional genes on blood derived B cells.

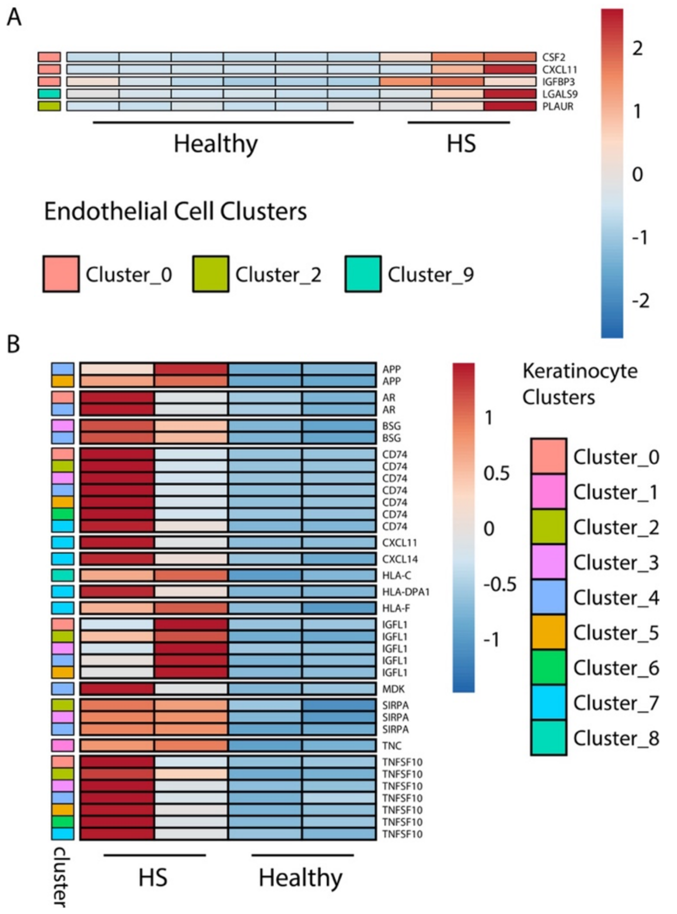


**Fig S4.** (A) Row-normalized heatmap depicting pseudo-bulk scRNASeq counts of significantly upregulated (adjusted p<0.05) genes in HS endothelial clusters versus healthy skin. (B) Row-normalized heatmap depicting pseudo-bulk scRNASeq counts of significantly upregulated (adjusted p<0.05) genes in HS keratinocyte clusters versus healthy skin.
